# Phenotyping women based on dietary macronutrients, physical activity and body weight using machine-learning tools

**DOI:** 10.1101/587220

**Authors:** Ramyaa Ramyaa, Omid Hosseini, Giri P Krishnan, Sridevi Krishnan

## Abstract

**Background:** Nutritional phenotyping is a promising approach to achieve personalized nutrition. While conventional statistical approaches haven’t enabled personalizing well yet, machine-learning tools may offer solutions that haven’t been evaluated yet.

**Objective:** The primary aim of this study was to use energy balance components – input (dietary energy intake and macronutrient composition), output (physical activity) to predict energy stores (body weight) as a way to evaluate their ability to identify potential phenotypes based on these parameters.

**Methods:** We obtained data from the Women’s Health Initiative –Observational Study (WHI-OS) from BioLINCC. We chose dietary macronutrients – carbohydrate, protein, fats, fiber, sugars & physical activity variables – energy expended from mild, moderate and vigorous intensity activity h/wk to predict current body weight either numerically (as kg of body weight) or categorically (as BMI categories). Several machine-learning tools were used for this prediction – k-nearest neighbors (kNN), decision trees, neural networks (NN), Support Vector Machine (SVM) regressions and Random Forest. Further, predictive ability was refined using cluster analysis, in an effort to identify putative phenotypes.

**Results:** For the numerical predictions, kNN performed best (Mean Approximate Error (MAE) of 2.71kg, R^2^ of 0.92, Root mean square error (RMSE) of 4.96kg). For categorical prediction, ensemble trees (with nearest neighbor learner) performed best (93.8% accuracy). K-means cluster analysis identified 11 clusters suggestive of phenotypes, based on significantly improved predictive accuracy. Within clusters, individual macronutrient gain and loss modeling identified that some clusters were strongly predicted by dietary carbohydrate while others by dietary fat.

**Conclusions:** Machine-learning tools in nutritional epidemiology could be used to identify putative phenotypes.

## Introduction

Nutritional phenotyping, a way to classify individuals based on their nutritional and health status, is a primary way to personalize nutrition recommendations [1]. Several approaches to phenotyping using metabolomics, nutrigenomics, metabonomics, epigenomics, and bioinformatics have come into existence [2]. However, there is yet to be a consensus as to how to bridge these phenotyping tools to personalize nutrition.

Within the framework of these phenotyping tools, data analyses play a significant role [3]. Statistical prediction models have been used in nutrition as early as the 1900s. The earliest and simplest examples based on classical linear regression analyses such as Harris-Benedict equation[4], or the Weir equation[5] are well known tools used in energy metabolism. These regression models ascribe to the one-size-fits-all theme that has been largely used in nutrition science. In recent years, as the dimensionality of data increased, as more variables were being measured, more complex tools became necessary to build predictive models. And personalizing efforts using more complex statistical analysis are becoming more common [6]

Traditional multivariate tools such as principal components analyses or hierarchical cluster analyses enable grouping people into categories, and are ideal to identify metabolic or nutritional phenotypes, and have been used to this effect in the past [7]. Linear and other discriminant analyses exist that do provide predictive capabilities. However, machine-learning tools appear to offer promising resources that can simultaneously create clusters and predict within the models that are built [8, 9]. The provision of validated predictive modalities is crucial to bridge the gap between phenotyping and customizing health solutions [10]. Furthermore, machine learning tools can develop models without pre-defined structure, local models (multiple models with each model capturing a locality of the multi-dimensional data space, as opposed to traditional global models which attempt to capture the entire multi-dimensional space of the data in one model), ensemble models (multiple models capturing different subsets of the data with no restriction of locality) etc. Ensemble models and local models are particularly useful when the data contains many sub-groups, since each sub-group can be modeled by one of the several models developed.

It is possible to train a machine learning based model on a wide multi-omic dataset to predict outcome measures, and has successfully been used by Zeevi et al to address changes in post prandial glycemic response (PPGR)[11]. A different approach is undertaken here. Our objective was to be able to identify sub-populations or ‘phenotypes’ within a large population of women, based on the relationship between body weight and dietary macronutrients/physical activity/socio-economic variables. To this end, we trained machine-learning algorithms on the macronutrient composition of diets, physical activity as well as other pertinent demographic data, to predict current body weight. Our goals were understanding (a) which commonly used algorithms achieve best prediction and why, (b) how these algorithms group women into clusters that might reflect underlying phenotypes, and finally (c) to evaluate how the relationship between dietary, physical activity and socio-demographic variables to body weight is different between these clusters. This paper presents several such models, and their evaluation to identify ideal prediction algorithms for this data, based on their ability to predict body weight both numerically, as well as categorically (BMI categories)

## METHODS

### Data Acquisition

We obtained data from BioLINCC, an online repository of epidemiological and clinical data hosted by the National Library of Medicine. We used the data from the Women’s Health Initiative-Observational Study (WHI-OS). This is a long-term national health study on postmenopausal women, which was started in 1991. The primary aim of this study was to identify strategies to prevent breast and colorectal cancer, heart disease and osteoporosis in postmenopausal women. There were two components that were initially started as part of the WHI - the randomized controlled Clinical Trial (CT) and the Observational Study (OS). The WHI-OS was aimed at observing and analyzing how well lifestyle behaviors such as diet, exercise and prior disease risk factors predicted disease outcomes, primarily heart disease and cancer. Enrolment for WHI-OS started in 1994, and was completed in 1998. A total of 93,676 women were recruited for this study. The selection criteria for recruitment was that the women were postmenopausal, between 50-79 y of age, ability and willingness to provide the information at baseline and follow up visits, and were planning to reside in that area for a minimum of 3 years. For the sake of answering our primary question, we used the WHI-OS data. Data that is of interest to our question was available from baseline. As part of the WHI-OS, women answered a modified semi-quantitative Block food frequency questionnaire in 1994 about their dietary intake between the years 1993-1994. In addition, they also filled out a physical activity questionnaire that categorized their activity levels into mild, moderate and vigorous physical activity [12]. Further, we obtained information about their health status such as presence of diabetes or hypertension, and demographic information such as age, ethnicity, education, income and marital status. Education and income were combined to arrive at one socio-economic score based on the method devised by Green LW, 1970 [13]. UC Davis IRB exempted review and approved the use of this data. All data treatment and analyses were done in Microsoft Office Excel (Redmond WA) JMP Pro 14.1 (Cary NC), R version 3.1.1[14], Matlab version 2017a

### Data cleanup

If total energy intake was reported to be <500 Kcal/d or >3500 Kcal per day, they were considered outliers and removed from the dataset. We only considered data where energy intake per kg body weight was between 15-35 kcals. Mahalonobis distance was used to evaluate outliers. These outliers were removed from the dataset to ensure robust model development. In the final analysis, data from 37,601 subjects were used to train the algorithm (“known data”), to predict the weight of 3706 “test” subjects. Figure 1 depicts the process of final study volunteer n identification.

**Figure 1:**
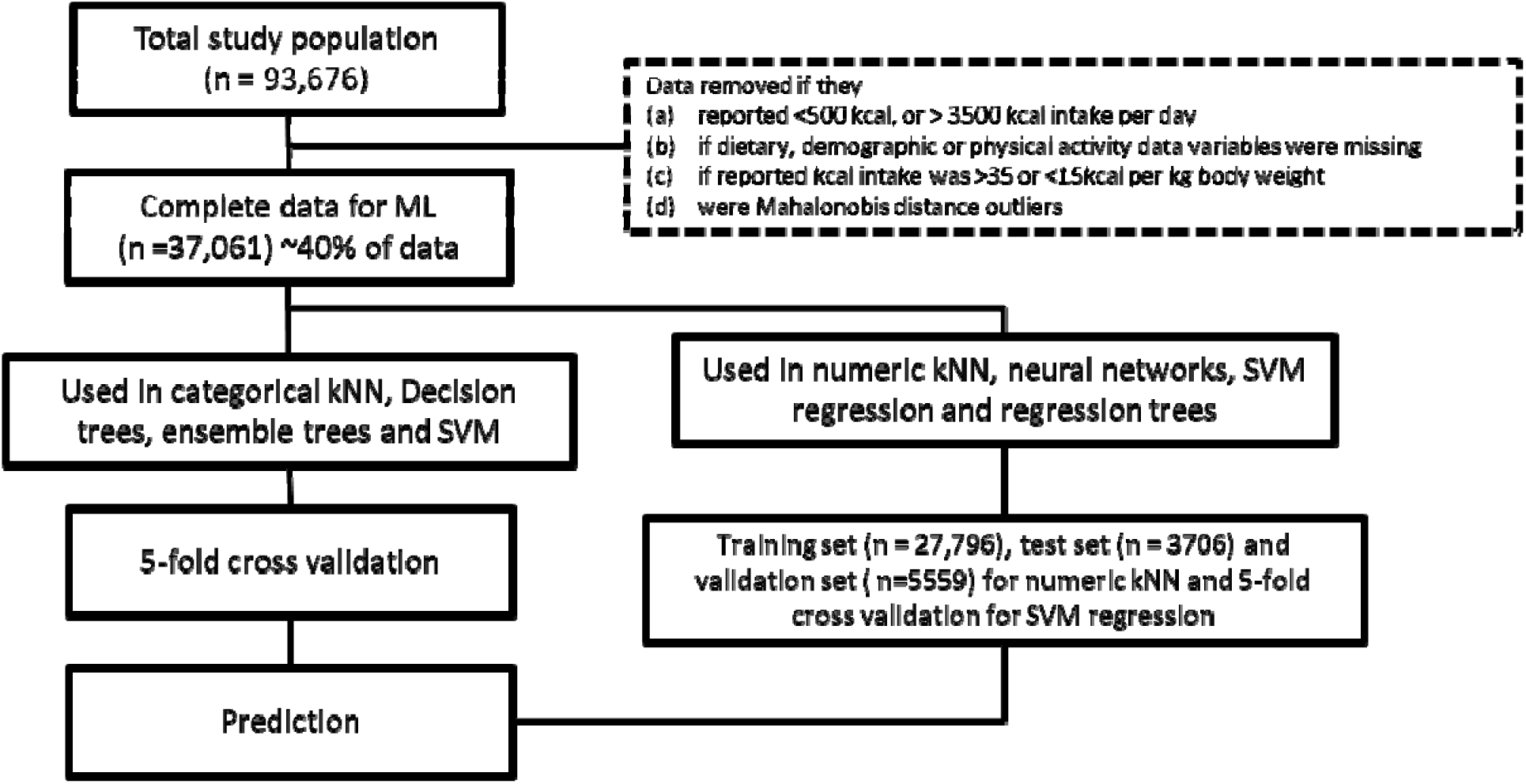
Flow-chart presenting data cleaning and prep process, including machine learning (ML) algorithms applied. kNN - k nearest neighbors, SVM - support vector machine.

### Feature selection

Dietary data collection methods, its implementation and the obtained data variables from the WHI-OS study are discussed here [12]. An extensive feature selection process was used that included standard least square regression, stepwise multiple linear regression, PLS regression and variable cluster analyses that returned a total of 29 variables with a p<0.0001 significance of being associated with body weight. For this first pass approach, 140 dietary variables, 12 anthropometric variables and 3 physical activity variables, as well as chronic disease state information were used. However, since our objective was to use macronutrients that will be pertinent to body weight, from amongst these selected features, only variables of physiological relevance were chosen, such as total energy (kcals), dietary fat, dietary protein, dietary carbohydrates, dietary sugars and dietary fiber to represent their diet. In addition, mild, moderate and vigorous intensity physical activity, height, socio-economic score and marital status were retained in the final model. All the disease states that were significant in this feature selection process were also retained in the final model.

### Data Preprocessing

Normalization: The input features were in various units with mean values spanning orders of magnitude. Many ML algorithms are very sensitive to differences in the scale of magnitudes of different inputs. To address this we normalized the data. We used a linear scaling to normalize the data to be in the interval [0,1].

### Principal Component Analysis

Principal component analysis (PCA) converts a set of observations of correlated variables into a set of values of linearly uncorrelated variables called principal components. Using an orthogonal transformation does this. The number of principal components is less than or equal to the smaller of the number of original attributes. This transformation is done in a way that the first principal component has the largest possible variance, and each succeeding component, in turn, has the highest variance possible under the constraint that it is orthogonal to the preceding components. We did PCA on our input features as a way of dimensionality reduction, and increase independence among inputs to the machine learning algorithms,

## Numerical Prediction

### Predictive modeling using machine-learning algorithms

For numerical prediction, the numerical value of weight in kg was predicted from the input variables. Many ML methods were tried – simpler ones such as statistical regression, regression tree, did not yield good results (data not shown); neural networks and SVM (explained below) are powerful techniques, which are capable of learning complicated functions; the method that worked best for this data is a local, instance based learning method called k-NN (explained below).

#### Regression SVM

Support vector machines or support vector networks are supervised learning algorithms and are commonly used when input vectors are non-linearly mapped to a very high dimensional feature space, which has a linear relationship to the output. This ensures high generalizability of the learning machine [17]. The algorithm uses kernels function to measure the similarities between two different patterns by returning a real value. One simple but not sufficiently general way to measure similarity in kernel function is using canonical dot product [17]. Vapnik (1995) presented *ε - insensitive loss function* to compute SVM regression [21].

This method is very powerful for non-linear analysis, but the search space in which to find the non-linear mapping becomes too large for high dimensional input data, and poses a practical challenge in using this [19,20].

#### Neural Network

Neural networks are biologically inspired and consist of a large number of very simple and independent processing units (neurons) which are connected unidirectionally. Neural networks are a powerful tool to capture the semantics or dynamics of very nonlinearly dependent factors. The first neural network model designed by McCulloch and Pitts in 1946 [20]. Rosenblatt introduced the first perceptron model and discussed its convergence to correct weights [23, 24, 25, 26]. Parkers and Rumelhart et al, introduced the back-propagation multilayer’s Neural Network model for weights determination [27, 28]. The back-propagation Marquardt algorithm for nonlinear least squares uses feed-forward neural networks for training. Compared with a conjugate gradient and a variable learning rate algorithms, the algorithm Marquardt is much more efficient when the network contains no more than a few hundred weights and also in many cases, the Marquardt algorithm converges while the mentioned algorithms fail to converge [29, 30, 31]. For this reason, in the present study, the Marquardt algorithm was chosen.

### kNN (k Nearest Neighbors)

Instance-based learning approaches such as kNN theoretically follow the straightforward way to approximate real or discrete valued target functions [15, 16]. The learning process is lazy and consists of storing training data; predicting the output of a new input vector involves fetching similar instances from saved training data and aggregating their outputs. Unlike many other techniques that build only one local approximation to the target function, one significant advantage of instance-based algorithms is that for each new query instance the model can build a new approximation to the target function. This gives instance-based algorithms, specifically case-based algorithms, the ability to capture very complicated relationships between attributes and outcomes There are two big disadvantages associated with the instance-based approach: (i) The cost of classification is much higher than other methods, since all computations are performed at the classification time rather than while training; (ii) This method incorporates all attributes of the instances, when the algorithm tries to bring back similar training examples from the memory. If the target variable is only dependent on few of the attributes, then this can cause instances that are very similar to be predicted further apart with a large distance [17, 18].

In k-nearest neighbors’ method, which is the most basic algorithm among instance-based methods, all instances are mapped to points in the n-dimensional space (ℛ^*n*^). Different distance measurement techniques can be applied to calculate nearest neighbors. The original algorithm uses standard Euclidean distance method. It should be noted that Euclidean distance and square Euclidean distance are usually used when data is not normalized. These two methods are also very sensitive to the scale of different independent attributes and having one or more attributes with big scale can decrease the effect of other attributes [19]. The City Block (Manhattan) distance between two independent attributes, unlike the Euclidean distance, is measured as the distance along x-axis plus y-axis. For numeric kNN method 75% of the dataset was assigned to training subset and the remaining 25% to testing. To validate the models, 5-fold cross-validation was used.

#### Information gain and loss models

In order to study the effect of each independent variable on the outcome accuracy, all independent variables were excluded from the model one by one and a new model was developed based on the other independent variables. Three different sets of models were observed based on the effect of variable exclusion, a) Removing variables improved the accuracy, b) Removing variables had no specific effect on outcome accuracy, and c) Removing all variables in part *<u>a</u>* and *<u>b</u>*.

### Categorical prediction

Here, independent attributes were used to predict BMI categories (underweight (*BMI* < 18.5), normal weight (18.5 ≤ *BMI* < 25), overweight (25 ≤ *BMI* < 30), grade I obese (30 ≤ *BMI* < 35), grade II obese (35 ≤ *BMI* < 40) and grade III obese (40 ≤ *BMI*)). For categorical approach Bagged Tree, Decision trees, SVM, kNN and Ensemble trees were evaluated (as before we are not presenting methods that were not promising).

### Decision Tree

Decision tree learning algorithms are one of the most effective and widely used inductive inference methods for discrete valued target functions [15]. A decision tree is tree-structure of Boolean questions about the input variables, with each branch ending in a leaf marked by an output category. If the input variables are such that a particular branch would be traversed, the corresponding leaf is the predicted classification of BMI.

Decision tree learning algorithm is a top-down, greedy algorithm, which constructs the tree as follows: Question the input attribute that has the most mutual information with the output, and ask the question that will reduce the information content (entropy) of the output; repeat this for each branch corresponding to each answer of the question. The ID3 decision tree algorithm was used for this dataset.

### Ensemble methods

In order to improve the generalizability and robustness of a predictor, results obtained from several basic predictors of a given learning algorithm can be combined. Ensemble methods try to combine the obtained results to achieve such goal. Ensemble methods can be categorized into two general groups. a) averaging methods such as bagging methods or forests of randomized trees, b) boosting methods such as AdaBoost or gradient tree boosting [20].

Bootstrap aggregating (bagging), is a machine learning ensemble meta-algorithm designed to improve the stability and accuracy of machine learning algorithms such as decision trees, neural networks, and linear regression. Generally, bagging predictor is a method of using multiple versions of base predictors to obtain an aggregated predictor. When the outcome is discrete-valued, the aggregation uses averaging over all different versions while for the categorical outcome it uses plurality vote to obtain the best accuracy. One of the main problems related to prediction methods is instability. Bagged methods could increase the stability when altering the learning set makes a huge difference in the constructed predictor [21]. The major difference between bagging methods arises from how the random subsets of training data are chosen. Some of these methods are pasting, bagging, random subspace, and random subspace [20].

Boosting methods follow a different logic than bootstrap aggregating. Many weak models are combined to produce an ensemble model that is more powerful. The base predictors are generated sequentially and one tries to reduce the bias of the combined predictor [22]. One of the most popular boosting algorithms is AdaBoost, which was introduced by Freund and Schapire in 1995 [23]. The algorithm uses many weak learners that just perform/predict slightly better than random guessing. Following this, it fits these weak learners repeatedly on the data and then uses weighted majority vote or sum technique to combine the results for the final prediction.

### Validation

For all machine-learning methods, there is always the risk of over-fitting or underfitting. Over-fitting happens when the model is too complex for the data and it is due to the small size of the dataset or presence of too much noise in the data. In the case of overfitting, the complex generated model captures the noise in training dataset. When overfitting happens the algorithms show low bias (error) and high variance. In contrast, under-fitting occurs when the statistical model or machine-learning algorithm cannot capture the underlying trend of the data and consequently the model does not fit the data correctly. In case of under-fitting occurrence, high bias and low variance are obvious.

Both under-fitting and over-fitting have problematic consequences for machine learning algorithms and lead to poor predictions for unseen input instances. The validation methods are essential tools in machine learning algorithm to make sure the constructed model is suffering from neither underfitting nor overfitting. A very basic approach called validation set method is randomly dividing the dataset into training and testing (70% training and 30% testing) subsets and build the model based on training data and then test the model’s performance with the testing subset. While this method is very fast and easy to implement, it also has some drawbacks. Test error can be highly variable depending on which observations are included in testing and training datasets. The other problem related to validation set method is that the model is developed on only a subset of the data and this potentially can lead to higher estimation error and consequently poorer model in the test phase.

To address the problems related to the validation set method, cross-validation tries to create a test dataset during the training phase by partitioning the dataset into subsets and using one subset called validation dataset for testing and the rest for training purpose. There are different cross-validation methods such as leave one out (L-O-O) and k-fold validation. In L-O-O the model is being trained on *n* – 1 observations and validated on the single validation set observation. Test error based on only one single observation is highly variable, however if the process get repeated for all instances in the dataset, then the average of all these test errors gives the overal error of the dataset. Having less bias in regression coefficients and no parameter estimation’s variation across the tarianing dataset are two big advantages of this method, however for large datasets it is computationally expensive.

The k-fold cross validation method is a compromise between two mentioned methods, which was used in the current manuscript. The dataset is randomly divided into subsets (aka fold). One subset is being used to test the model and the other *k* − 1 subset train the model. This process is repeated *k* times for all subsets and the average of the these *k* test errors presents the overal error of the dataset. This method requires less computational resources, the estimation is more accurate than L-O-O (L-O-O has less bias than k-fold cross validation but larger variance) [20].

### Clustering

Powerful ML methods such as NN and SVM can fail for many reasons including the presence of many dimensions, potentially irrelevant ones, too much noise etc. For our dataset, the training set error was quite high, indicating that the desired function was not being learned. This coupled with the fact that local and ensemble models perform well led us conjecture that for similar values of inputs, we had widely varying outputs. In such cases, there may be hidden variables which determine the outputs, or the function from inputs to outputs. If this were the case, clustering would improve the performance of the algorithms.

Cluster Analysis groups a set of objects that have the maximum similarities. Hierarchical clustering analysis (HCA) or connectivity based clustering follows the simple idea that the nearby objects are more related to each other than the ones further away. HCA, which is a greedy algorithm, falls in two different categories, Agglomerative (Linkage) and Divisive (K-means and K-medoids). In agglomerative (bottom-up) clustering, each object starts in its own cluster and a pair of objects can merge as one moves up, however divisive cluster analysis which is a top-down method, all objects are in one cluster and different clusters dis-join recursively as one moves down.

K-Means is the simplest and most commonly used algorithm which uses squared error criterion. This algorithm represents clusters by their centroids, which is the mean of all objects in one specific cluster. The algorithm for K-means clustering uses partitioning error to stop the iteration. This algorithm can be presented as a gradient-descent procedure, which starts with an initial fixed number of cluster centroids and constantly updates them to reduce the error as much as possible. K-means algorithm has a linear time complexity and this makes the algorithm popular among researches with large data sets[24].

Clustering algorithms’ performance measures quantify the proximity of the points in each cluster, and the distance separating points from different clusters. There are several measures, and they all measure the distance between each data point from the centroid of its cluster (as a measure of cluster cohesiveness) and the distance between centroids of various clusters (as a measure of cluster separation), and combine the two measures. The clustering algorithms we use were sensitive to (a) number of clusters which is a parameter specified by the user (b) initial random clustering.

The clustering algorithms are very fast, and so are run many times with may different initialization for each desired cluster number, and a maximum value of the desired criteria of inter-cluster separation and intra-cluster cohesion is chosen. This process repeated varying the number of clusters to determine the ideal number of clusters. The Calinski-Harabasz criteria was used in the current study to determine the goodness of clustering [25]. The higher the score, the better the clustering, between 1 − 15 cluster numbers were evaluated for this study, and the cluster number corresponding to the highest criteria score was chosen.

## RESULTS

The independent variables that were used to predict body weight as well as BMI category are listed in Table 1.

**Table 1:** Feature selection based final list of variables chosen for building models.

| **Independent variables** | **Variable type** |
| --- | --- |
| Energy (kcal) | Numeric |
| Carbohydrates (g) | Numeric |
| Sugars (g) | Numeric |
| Protein (g) | Numeric |
| Fat (g) | Numeric |
| Fiber (g) | Numeric |
| Age (y) | Numeric |
| Ethnicity | Categorical |
| Socio-Economic Score | Numeric |
| Marital status | Categorical |
| Vigorous intensity physical activity (h/wk) | Numeric |
| Moderate intensity physical activity (h/wk) | Numeric |
| Mild intensity physical activity (h/wk) | Numeric |
| Height (cms) | Numeric |
| Overactivethyroid | Binary |
| Underactivethyroid | Binary |
| Heartfailure | Binary |
| Angina | Binary |
| Atrialfibrillation | Binary |
| Kidneyorbladderstones | Binary |
| Dialysisforkidneyfailure | Binary |
| Stomachorduodelulcer | Binary |
| Diverticulitis | Binary |
| Pancreatitis | Binary |
| Liverdisease | Binary |
| Alzheimersdisease | Binary |
| Multiplesclerosis | Binary |
| Parkinsonsdisease | Binary |
| ALSLouGehrigsdisease | Binary |
| **Dependent variables** |  |
| Body weight (kg) | Numerical |
| BMI (kg/m2) | Categorical |

### Numeric prediction models

Table 2 summarizes the results from different models evaluated for numerical prediction using the independent variables. Gaussian Regression SVM performed the worst (MAE 6.30 kg, and a *R*^2^ of 0.70) and kNN performed the best (MAE 2.71 kg, and *R*^2^ of 0.92). The two-layer feed forward neural network trained with the Leavenberg-Marquardt algorithm and 200 hidden neurons was not as successful at predicting body weight (MAE: 5.93kg, and *R*^2^ 0.73). For the kNN model, which was the best fit, figure 2 is a scatter plot that presents the relationship between predicted weights vs. real weights, as well as the distribution of error in prediction. As observed, despite being the best fit, the overlap between the real weights (black dots) vs. predicted weights (red dots) is not perfect, with several outliers.

**Table 2:**
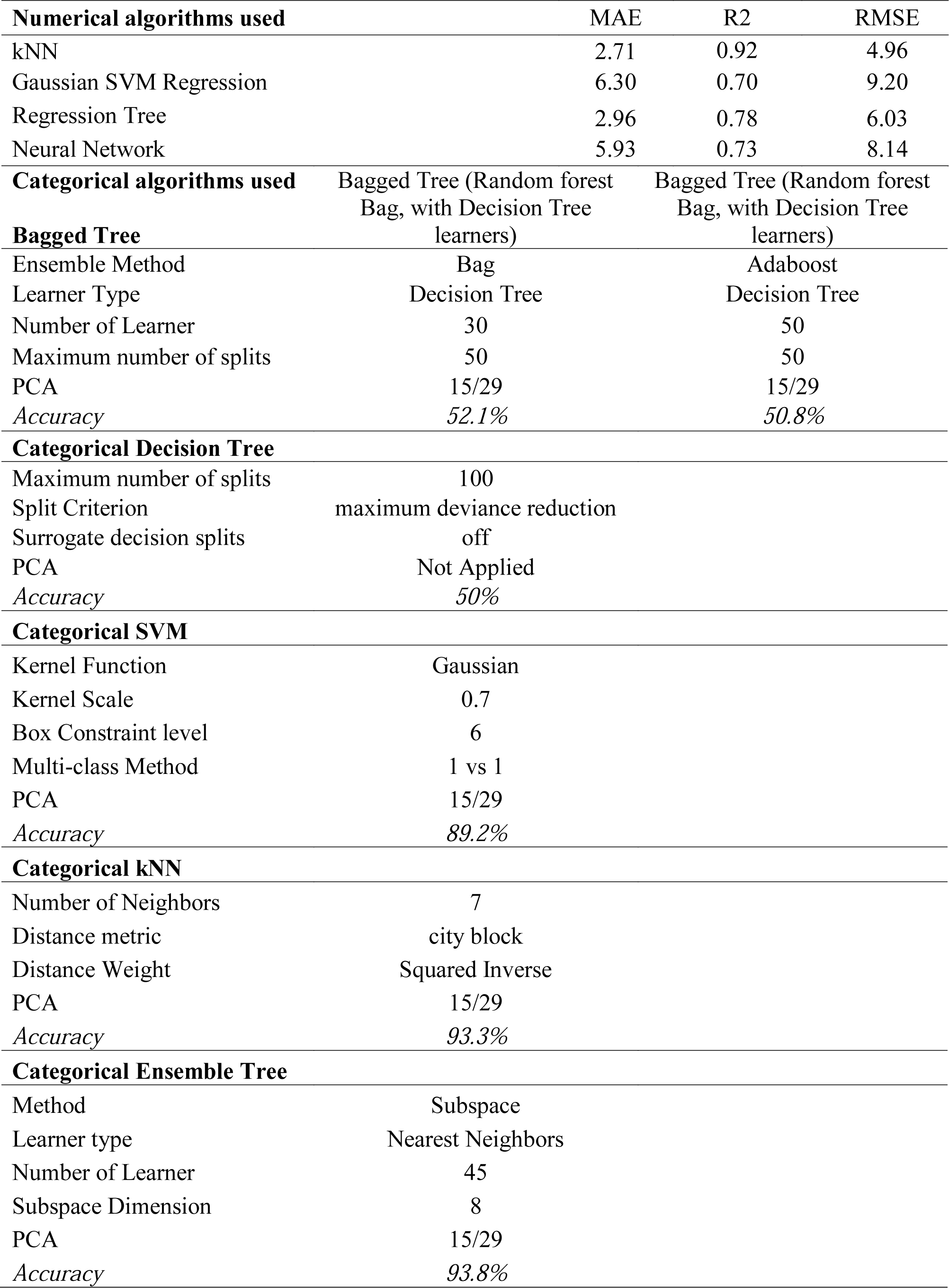
Numerical and categorical approach summaries

**Figure 2:**
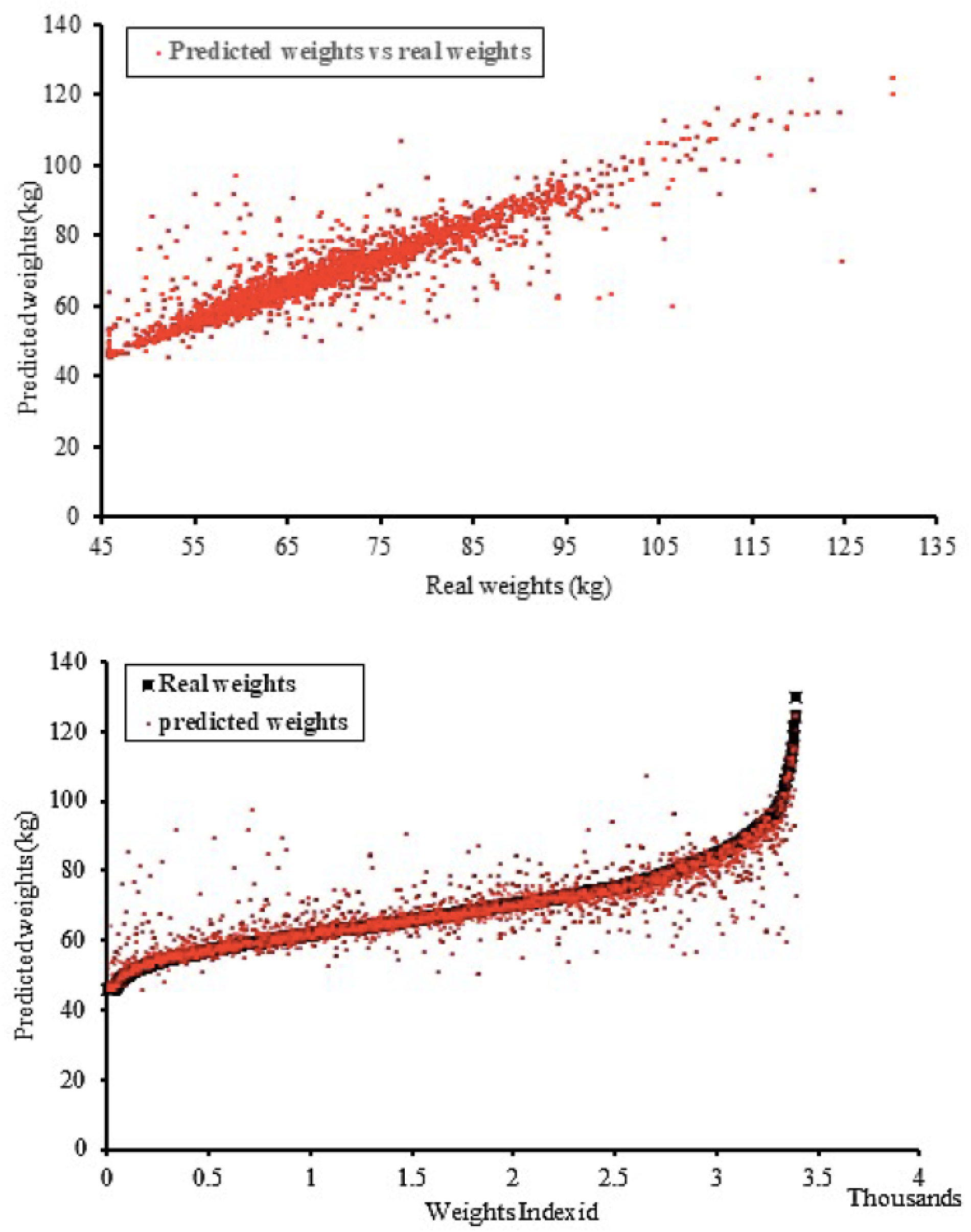
The top panel shows comparison between predicted weights by kNN vs. real weights, and the bottom panel shows a scatter plot comparing predicted and real weights vs. weight index. In the bottom panel, the red dots are the predicted weights and the black squares are the real weights

### Information loss and gain model evaluations

We evaluated the effect of elimination of each independent variable on MAE, *R*^*2*^ and RMSE (Table 2). Removing different variables had a different impact on the output accuracy as reported, and removing energy (kcal) and moderate intensity physical activity (h/wk) improved the accuracy the most (MAE 1.76 kg, and a *R*^*2*^ *of* 0.94). Socio-economic score, mild physical activity, age, ethnicity, carbohydrates and fiber, were identified as ineffective when removed individually to the overall model outcome. However, removing all of them at once resulted in a changed model with the highest error (MAE 3.27 kg, and a *R*^*2*^ *of* 0.90). It is interesting to note that removing fat worsened the model fit (MAE 2.83 kg, and a *R*^*2*^ *of* 0.91), while removing dietary carbohydrates did not affect the model significantly (MAE 2.73 kg, and a *R*^*2*^ *of* 0.92).

### Categorical prediction models

Ensemble trees using nearest neighbors as the learner type were best able to predict BMI category (93.8% accuracy), closely followed by actual kNN (93.3% accuracy), while decision tree models had the worst fit (50% accuracy). This is depicted in Table 2, along with the implementation details for the model. Figure 3 displays the confusion matrix outlining the positive predictive value as well as the false discovery rate for the ensemble tree model with the highest accuracy. The normal weight BMI category had the highest positive predictive value, while underweight as well as grade I obese categories had the lowest. The highest misclassifications were also in these categories − 7% of grade I obese were misclassified as grade II obese, and 12% of normal weight were misclassified as underweight.

**Figure 3:**
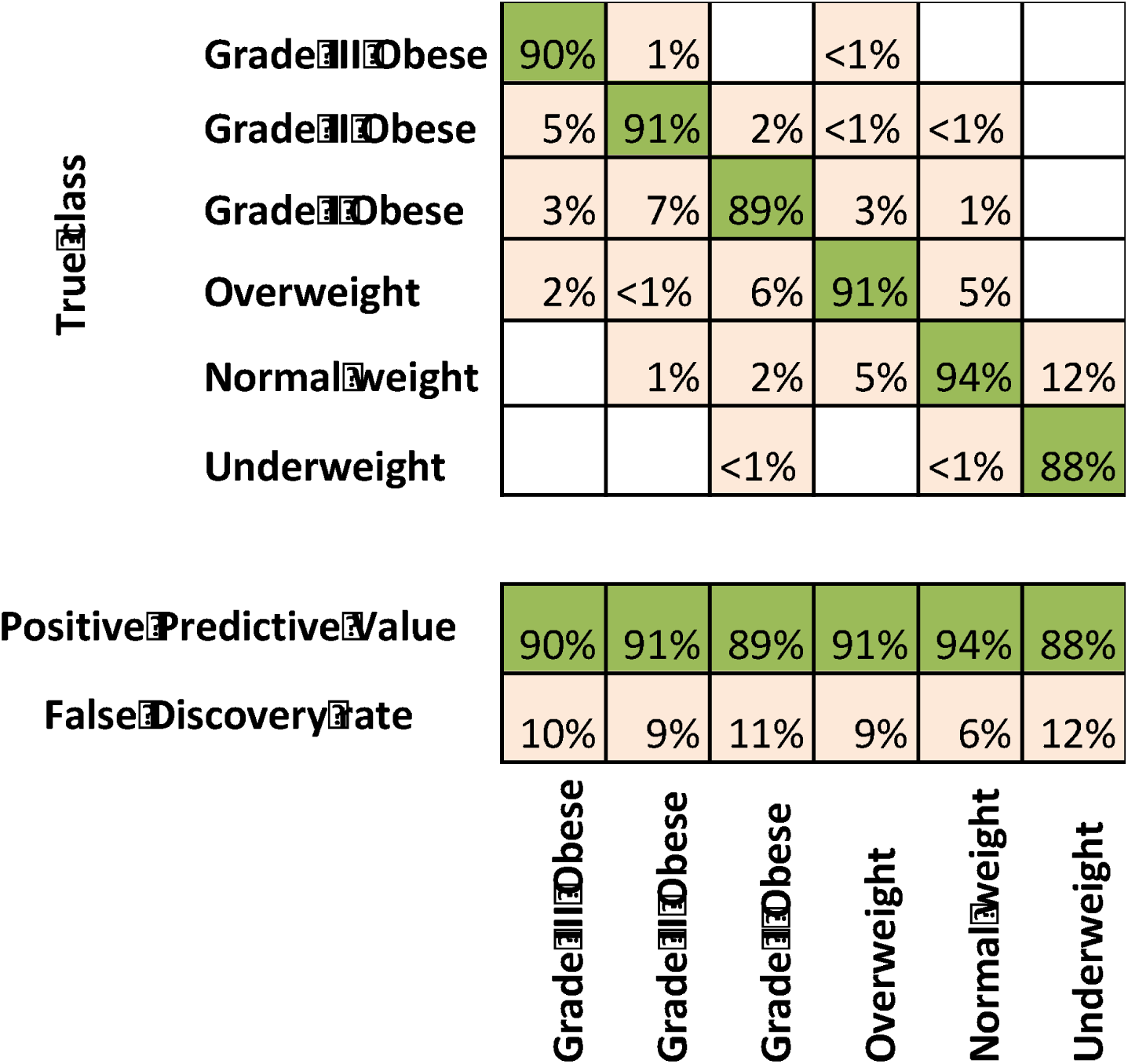
Confusion matrix for the Ensemble Tree with subspace method using nearest neighbors’ learner. Green color boxes indicate positive prediction and cream color boxes indicate false predictions.

### Cluster analysis

While global models did not successfully predict the outcome variable, local models performed a little better. However, even those were not able to predict to the closest kg body weight as could be expected. So, cluster analyses were done to see if they improve model fit and prediction. Since local models were more successful than global models, a local unsupervised learning clustering tool (K-means clustering) was used to identify ‘phenotypes’ within this population. K-means cluster analysis identified 11 clusters of women.

A summary of their fit characteristics is given in table 4. It is important to note that once clusters were formed, kNN performed significantly better within each cluster at predicting body weight using the same input variables, than using the population as a whole. However, the R^2^ value was lower for the clusters compared to the whole dataset. R^2^ is the ratio of the variance in the dependent variable and the proportion of variance that is explained by the independent variables. A reduced R^2^, but improved predictability (reduced MAE and RMSE) suggests that a lesser proportion of variance in the dependent variable is explained by the independent variables, but there is overall lesser variance to predict, so the predictability does not suffer.

**Table 3.** Further exploration of the best-fit kNN model

| *Variable* | *MAE* | *RMSE* | *R<sup>2</sup>* | **Exclusion Effect** |
| --- | --- | --- | --- | --- |
| **With all Independent Variables** | **2.71** | **4.96** | **0.92** |  |
| Without energy (kcal) | 1.77 | 4.30 | 0.94 | Improve |
| Without moderate intensity physical activity (h/wk) | 2.69 | 4.79 | 0.92 | Improve |
| Without dietary protein (g) | 2.77 | 5.04 | 0.92 | Reduce |
| Without marital status | 2.81 | 5.10 | 0.91 | Reduce |
| Without total fat (g) | 2.83 | 5.19 | 0.91 | Reduce |
| Without dietary sugars (g) | 2.81 | 5.04 | 0.92 | Reduce |
| Without vigorous intensity physical activity (h/wk) | 2.78 | 5.02 | 0.92 | Reduce |
| Without height (cms) | 3.04 | 5.30 | 0.91 | Reduce |
| Without Socio-Economic Score | 2.74 | 4.90 | 0.92 | No effect |
| Without mild intensity physical activity (h/wk) | 2.78 | 4.96 | 0.92 | No effect |
| Without age | 2.80 | 4.93 | 0.92 | No effect |
| Without Ethnicity | 2.71 | 4.93 | 0.92 | No effect |
| Without Dietary Total Carbohydrate (g) | 2.73 | 4.94 | 0.92 | No effect |
| Without Dietary Fiber (g) | 2.76 | 4.93 | 0.92 | No effect |
| **Without energy (kcal) & Moderate intensity physical activity (h/wk)** | **1.76** | **4.16** | **0.94** | **Improve** |
| Without all ineffective variables | 3.27 | 5.39 | 0.90 | Reduce |
| Without energy (kcal), moderate intensity physical activity (h/wk) and ineffective variables | 2.26 | 4.64 | 0.93 | Reduce |

**Table 4:** Clusters and their performance relative to the fit of complete data model

| Cluster number | Average body weight |  |  | Cluster size (n) | MAE | RMSE | R<sup>2</sup> |
| --- | --- | --- | --- | --- | --- | --- | --- |
|  | (kg) |  |  |  |  |  |  |
|  | Mean ± SD |  |  |  |  |  |  |
| All clusters | 81.9 | ± | 2.2 | 37,061 | 2.71 | 4.96 | 0.92 |
| Cluster number 1 | 85.0 | ± | 2.0 | 2169 | 0.49 | 1.10 | 0.70 |
| Cluster number 2 | 61.7 | ± | 1.7 | 5517 | 0.42 | 0.91 | 0.71 |
| Cluster number 3 | 78.5 | ± | 1.7 | 2763 | 0.58 | 1.32 | 0.69 |
| Cluster number 4 | 72.7 | ± | 1.6 | 4218 | 0.42 | 0.93 | 0.70 |
| Cluster number 5 | 123.8 | ± | 4.1 | 96 | 0.48 | 1.06 | 0.64 |
| Cluster number 6 | 67.4 | ± | 1.6 | 5336 | 0.37 | 0.88 | 0.70 |
| Cluster number 7 | 49.6 | ± | 2.3 | 1829 | 0.06 | 0.28 | 0.99 |
| Cluster number 8 | 102.2 | ± | 2.8 | 488 | 0.39 | 0.87 | 0.71 |
| Cluster number 9 | 111.4 | ± | 3.0 | 255 | 0.48 | 1.20 | 0.74 |
| Cluster number 10 | 56.0 | ± | 1.7 | 3894 | 0.55 | 1.34 | 0.77 |
| Cluster number 11 | 92.3 | ± | 2.4 | 1231 | 0.94 | 1.93 | 0.62 |

We used kNN and decision trees to perform information gain and loss evaluations. kNN identified that clusters 3, 11 and 9 improved fit (based on MAE, R^2^ and RMSE, data not shown) when dietary kcals, carbohydrates, sugars, protein, fat, fiber, or age at screening were removed from the models individually. But the other models worsened when these variables were removed individually. Height was the most important variable, and when removed, all clusters had poorer fit parameters (MAE, R^2^, RMSE) and the prediction worsened significantly.

Decision trees were used to determine which variables the algorithm chose to split first, to infer relative variable importance based on *information gain* as mentioned earlier. These results are given in Figure 4. The decision tree splitting process was stopped as soon as the first dietary macronutrient variable, and the first physical activity variables were identified to split by the algorithm. For clusters 1 and 8 (total n = 2657), dietary protein was a significant predictor of body weight, while for clusters 3 and 6 (n = 8099) it was dietary fiber. Dietary fat was pertinent in cluster 2 and 10 (n = 9411), sugar was a primary predictor in clusters 4 and 5 (n = 4314), while dietary carbohydrate was the pertinent predictor in clusters 7, 9 and 11 (n = 3315). While the global model suggested that dietary fat was an important predictor of body weight, the individual clusters appear to indicate a different story. Given that the predictive ability of these clusters are significantly better than that of the population model, it appears dietary carbohydrates (fiber, sugar as well) are primarily predicting body weight for a majority of the population. Five out of the 11 clusters indicated that mild-intensity physical activity was a primary predictor, while 2 for vigorous intensity physical activity, and the remaining 3 for moderate intensity physical activity. Total energy intake was the first split for 9 out of the 11 clusters, and height and age were the first split for the remaining 2 clusters. Spearman’s correlation analyses revealed significant associations between these macronutrients or macronutrient components and body weight within each cluster. For clusters 1, 2, 4, 5 and 8 the correlations match the decision tree’s prediction, but the correlations are less congruent for the remaining clusters.

**Figure 4:**
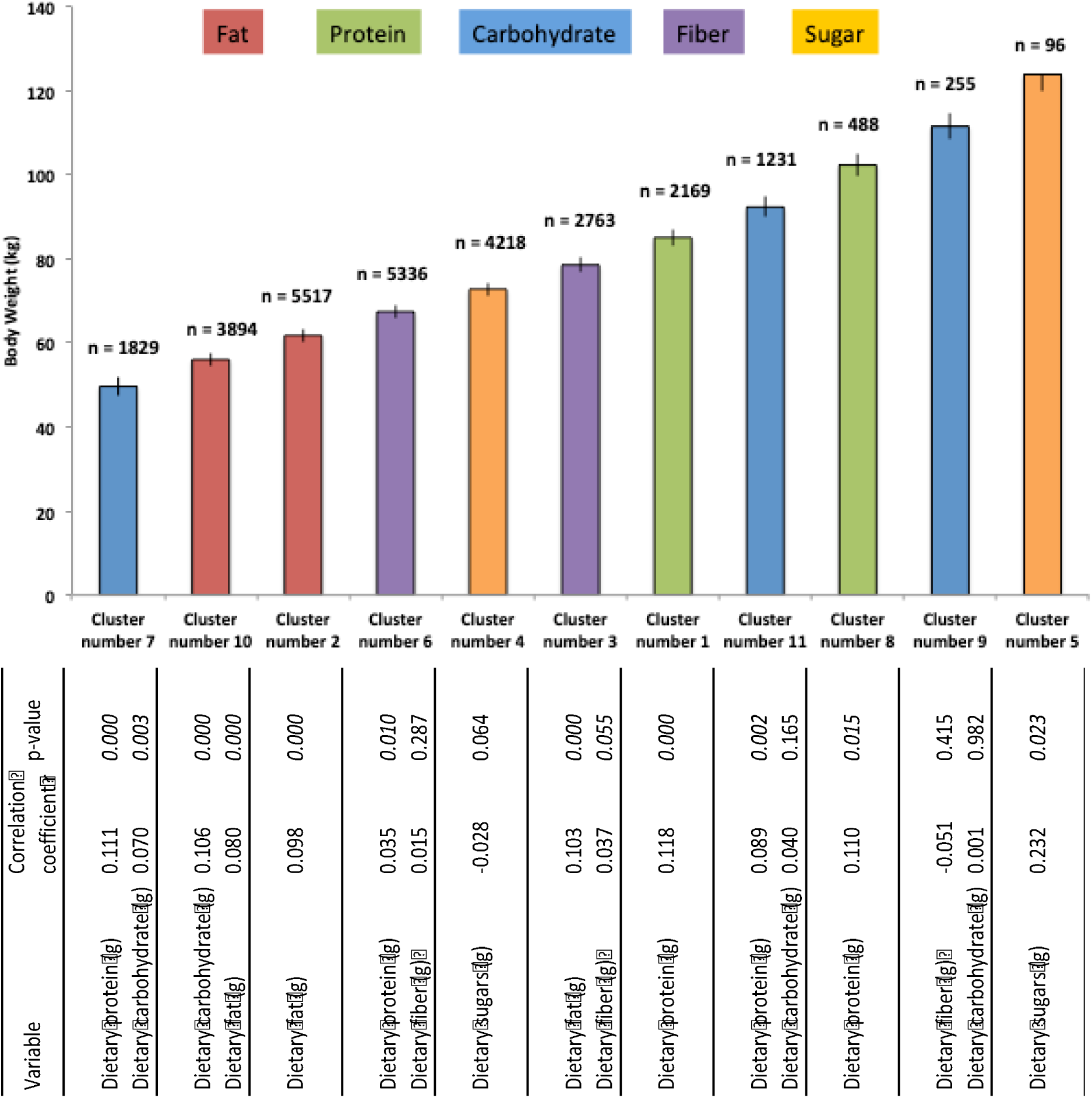
Decision tree splits for clusters. The figure shows the mean body weight for each cluster, and the n indicates the number of women per cluster. The colors identify which main dietary caloric or macronutrient component had the largest predictive ability for each cluster. Beneath the bar graph are the corresponding spearman’s correlation co-efficients (r) and p values (variables vs. body weight (kg)). If only one variable is listed, that was the largest r, as well as the lowest p-value. If two variables are listed, the first one is the largest r, and the lowest p-value, which does not match what the decision tree suggested. However, the second listing shows the r and p values for the variables picked by the decision tree.

## Discussion

To our knowledge, this is the first effort to explore the relationships between selfreported dietary macronutrient data and physical activity adjusting for socio-demographic and disease state variables to predict current body weight using machine-learning tools. Our effort was focused on using the ability of predictive modeling tools, as both predictive as well as an inferential tool. This study identified that the k-nearest neighbor algorithm was the best predictive tool for numerical prediction, and Ensemble Regression trees, also using nearest neighbor learning learner type, were the best for predicting BMI categories. The nearest neighbor approach appears best able to handle the noise present in self-reported dietary data.

While the overall data model showed reasonable fit and predictive ability, our clusters produced relatively superior fit statistics. The clustered data also suggested that for several clusters dietary carbohydrate in any form − complex carbohydrates, sugars or even dietary fiber were significant predictors of body weight (n = 15,728), and fat was significant for a lesser number of clusters (total n = 8099).

Parametric model fitting algorithms assume a predefined model template and model fitting involves determining the parameters of the model template that best fits the data [26]. For instance, linear regression assumes the model to have the template of a linear equation, and attempts to find the coefficients of the equation. Non-parametric models do not assume a predefined model, but instead attempt to derive both model structure as well as parameters from the data. These models are particularly useful when the independent variables do not individually determine the shape of the regression. Many machine-learning algorithms such as decision trees and neural networks are non-parametric (when the search for network structure is also included as part of model fitting).

Ensemble regression trees were our best-fit model algorithms to predict BMI categories. Local methods and Ensemble methods form special types of non-parametric model fitting. Ensemble methods do not develop a single global model. Instead they develop many models of a same template or family of templates. The ensemble of models is developed by a randomized method (the ensemble members are models that are randomly initialized, and upon learning, each member converges to a different local minimum based on its initial values). The predictions form these diverse models are then combined (in a way that increases accuracy) to form a final result. Empirically, ensembles tend to yield better results when there is significant diversity among the models so that each ensemble member captures a different subset of the data. Many ensemble methods, therefore, try to ensure that the ensemble members are diverse. One interpretation of the success of ensemble model is that each model used in the ensemble models a subset of the entire data set, and the combination of the models can be thought of as a global model.

For numerical prediction, kNN algorithm was our best fit. Local models such as kNN, similar to ensemble methods, do not develop a single, global model that fits the data; instead, they are a family of models around each data point such that a data point’s local model fits only the data in its local neighborhood. The main advantage of this method is that it can be used on data sets that would require a very complicated global model. The success of a local model, as seen in this report, implies that the data points close to each other form a model, which predicts the dependent variable well from the independent variables.

Further, in the setting of numerical predication, local modeling performed best. So, the locality of each point can be represented with a simple model. This opens the possibility that simple model templates have simple parametric phase spaces i.e., the locality of a data point in terms of the relationship between dependent and independent variables is controlled by one or more parameters, which may allow similarity measures, and consequently grouping. This possibility, along with the success of ensemble models suggests that it might be possible that the dataset may be grouped appropriately, and each group can be modeled using a single global model.

Ensemble models and local models performed the best for this data set. This suggests that the data set is too complex for a single complicated global model. Fitting a complicated model increases the possibility of over-fitting, as well as making parameter estimation challenging, not to mention the fact that it may not yield the best results. Local and ensemble models fitting the WHI-OS data best could be explained in many ways − such as the search space being too big. Alternately, hidden/latent variables (such as genetic information, other lifestyle characteristics, eating behaviors, or just errors in self-reports) may be hindering the ability of the global models to be successful. It is also very likely that there is a dynamical system involved indicating that our underlying assumption that body weight is a deterministic function only of diet, anthropometric and demographic characteristics is incorrect, and needs a time component to be successful.

The clustering efforts resulted in an improved prediction, suggesting that the high variance in the larger dataset was contributing to the larger prediction error. Further, the cluster analyses also identified sub-groups suggestive of phenotypes that are influenced by different dietary macronutrients. Dietary carbohydrates, or its sub-components appear to be predominantly influencing body weight in most of the clusters. Exceptions occur in few clusters where protein or fat are the predominant influences. Whether this suggests that the women in these clusters are more susceptible to these dietary components with respect to body weight gain is a potential hypothesis, which will need to be tested in future studies

One of the limitations of the current study is the use of self-reported dietary data. Mobile health apps, epidemiological studies as well as large health-tech companies do use such data, even though their accuracy is controversial [27, 28]. Self reported dietary data, especially in the WHI, has been criticized for its poor quality [29], with clear under-reporting (up to ∼20%, especially in the younger demographic), but unrelated to their BMI. This indicates that our evaluation of the relationship between diet and body weight/BMI are affected by these errors, since we studied all BMI classes as well as age groups. With respect to self-reported dietary data, the ease of collection, at such large scale is an advantage. This, when used along with application of robust data cleaning, pre-treatment and evaluation, as outlined in this report, can make self reported data effective. An alternate to using dietary self-reported data to represent nutrient intake, would be to use it to represent a dietary “pattern”, as is becoming more common recently [30]. This could be a promising future endeavor.

A recent review paper [31] suggested a framework for personalizing nutrition approaches. Our report is a first step in (a) evaluating a framework for personalizing nutrition using population data, and, (b) evaluating relationships between dietary caloric variables (macronutrients) and body weight suggestive of different phenotypes responding differently to dietary carbohydrates, fat or protein. This suggests that such frameworks/pipelines may be useful to personalize ideal dietary intake levels, beginning at the population level. Multi-center controlled feeding trials, providing appropriate diet to the appropriate clusters, will need to be done in order to evaluate the predictive ability of these models. Further, an ideal future follow-up project would be to use these tools to predict change in body weight using these variables, and similar approach. The promise of using machine learning tools to achieve nutritional phenotyping needs to be explored further to set up standard paradigms based on data-type, origin of data and the researchers’ hypothesis, among several other factors.

